# Natural variation in oxytocin receptor signaling causes widespread changes in brain transcription: a link to the natural killer gene complex

**DOI:** 10.1101/2023.10.26.564214

**Authors:** Arjen J. Boender, Zachary V. Johnson, Kayla K. Green, George W. Gruenhagen, Brianna E. Hegarty, Kengo Horie, Kiyoshi Inoue, Ewoud R.E. Schmidt, Jeffrey T. Streelman, Hasse Walum, Larry J. Young

## Abstract

Oxytocin (OXT) is a highly conserved neuropeptide that modulates social cognition, and genetic variation in its receptor gene (*Oxtr*) is linked to divergent social phenotypes. However, the molecular mechanisms connecting *Oxtr* genotype to behavioral outcomes remain obscure. Here, we leverage naturally occurring *Oxtr* polymorphisms in the prairie vole that associate with striatal-specific OXTR density to investigate how OXTR signaling influences brain function. Specifically, we identify OXTR-dependent transcriptomic changes in the natural killer gene complex (NKC) - a genomic region classically associated with peripheral immune function. Centrally, OXTR-regulated NKC genes are positioned to influence microglia-neuron interactions. Consistent with a role for these genes in shaping neuronal connectivity, we show that genetic reduction of OXTR levels leads to increased dendritic spine density on striatal *Oxtr*-expressing neurons. In addition, we provide support for a similar relation between variation in *OXTR* mRNA levels and NKC transcription in humans. Together, our findings suggest a role for OXTR signaling in the shaping of neural circuits through transcriptional control of the NKC, outlining a mechanism via which variation in OXTR signaling may influence circuit connectivity to generate diversity in social behaviors.

## Introduction

Oxytocin (OXT) is a highly conserved neuromodulator of social behaviors, and variations of this molecule have been identified in several invertebrates and most vertebrates^1^. This neuropeptide family has evolved from controlling gustatory plasticity and mating circuits to regulating a diverse range of complex social behaviors^2^. OXT primarily signals through its receptor (OXTR), which enhances the saliency of social stimuli^3^, and supports key aspects of social cognition, including social reward processing^4^, individual recognition^5^, social attachment^6^, and parental care^7^. Animal studies continue to elucidate neural circuits that are modulated by OXT and shape social phenotypes^8^, informing clinical studies to evaluate the OXT system as a target for the treatment of social disability, such as social anxiety or autism spectrum disorders (ASD)^9^. A detailed understanding of OXT signaling and its consequences on brain function will therefore provide valuable insight into the mechanisms that determine sociality in both health and disease.

The socially monogamous prairie vole (*Microtus ochrogaster*) shows notable similarities to human social behavior, as they console familiar conspecifics, form long-term pair bonds, and engage in biparental care^10^. Prairie voles express high levels of OXTR in the nucleus accumbens (NAC)^11^, a brain region that integrates sensory input with motivated actions^12^. OXT signaling in the NAC facilitates pair bond formation, as genetic or pharmacological manipulation of OXTR signaling in this area can facilitate or prevent mating-induced partner preference, a laboratory proxy for pair bond strength^13–15^. In contrast, promiscuous rodent species, such as the montane vole *(Microtus montanus)*, the house mouse (*Mus musculus*), and the Norway rat (*Rattus norvegicus*), show much lower levels of NAC OXTR^11,16^. A similar pattern is observed in primates^17^, as the polygamous rhesus macaque (*Macaca mulatta*) lacks NAC OXTR^18^, while the socially monogamous marmoset (*Callithrix jacchus*) expresses it abundantly^19^. Notably, humans also show high NAC *OXTR* expression^20^, consistent with a conserved role for NAC OXTR signaling in the evolution of social bonding.

Variation in NAC OXTR abundance is not only observed between species, but also within species, as prairie voles show considerable individual variation in NAC OXTR density. Compared to standard laboratory mouse strains, our laboratory prairie vole population exhibits greater genetic diversity. This feature previously enabled us to identify a set of single nucleotide polymorphisms (SNPs) in the prairie vole *Oxtr* gene locus that strongly predicts individual OXTR density in the NAC (*R*^2^>0.7), independent of sex and specific to the NAC^21^. Variation in NAC OXTR density not only affects correlated FOS activity directly after mating^22^, but is also associated with divergent performance in behavioral assays of sociality, such as partner preference^21^, social obervation^23^, alloparental care^24^, and resilience to neonatal neglect or paternal absence^25,26^. Moreover, NAC OXTR density correlates with divergent mating strategies in the wild^27^. Thus, variation in NAC OXTR signaling influences a range of social behaviors throughout development and may promote diversity in social strategies. Also in humans, SNPs in the *OXTR* gene predict its expression^28^, and associate with various aspects of sociality such as relationship quality^6^, resilience to childhood adversity^29,30^, and social recognition^31^. *OXTR* polymorphisms also predict brain function in relation to social disability^32–34^. Understanding the mechanistic link between *OXTR* variation and divergent sociality could offer critical insight into how social experiences shape behavior and may give clues to the complex etiologies of conditions of altered sociality, such as ASD and schizophrenia.

Building on our previously identified association between *Oxtr* genotype and NAC OXTR density, we here investigated how variation in OXTR signaling influences gene regulation and neurodevelopment. Using ATAC and RNA sequencing, we profiled genotype-associated differences in chromatin accessibility and gene expression. Although we did not detect differences in chromatin accessibility in the *Oxtr* gene locus, we found that *Oxtr* genotype predicts brain-wide transcriptional differences, beyond the NAC and *Oxtr* expressing cell types. Importantly, we demonstrate that many of these transcriptional changes are mediated by OXTR signaling. Notably, we show that OXTR signaling controls gene expression in the natural killer gene complex (NKC), a genetic region classically involved in innate immunity^35^, although some NKC genes have recently been implicated in neural development^36–38^. Indeed, we show that variation in OXTR levels affects dendritic spine density in naïve animals, supporting a role for OXTR signaling in the shaping of neural circuits during development. Our results give surprising clues as to how OXTR signaling shapes brain development, providing new hypotheses on the molecular origins of diversity in social behavior.

## Results

### *Oxtr* genetic variation associates with widespread transcriptional changes beyond the receptor gene itself

We previously demonstrated that NAC OXTR density in prairie voles is strongly predicted by a set of non-coding SNPs in the *Oxtr* gene locus (*R*^2^>0.7, Figure 1A-B)^21^. This variation in OXTR abundance is brain region-specific, as genotypes show dramatic differences in NAC OXTR density, but exhibit equal densities in the insular cortex (INS). These SNPs are in perfect linkage disequilibrium (LD), and we genotyped animals at position NW_004949099:26351196 to differentiate between low (T/T) and high (C/C) NAC *Oxtr* expressing animals.

**Figure 1.**
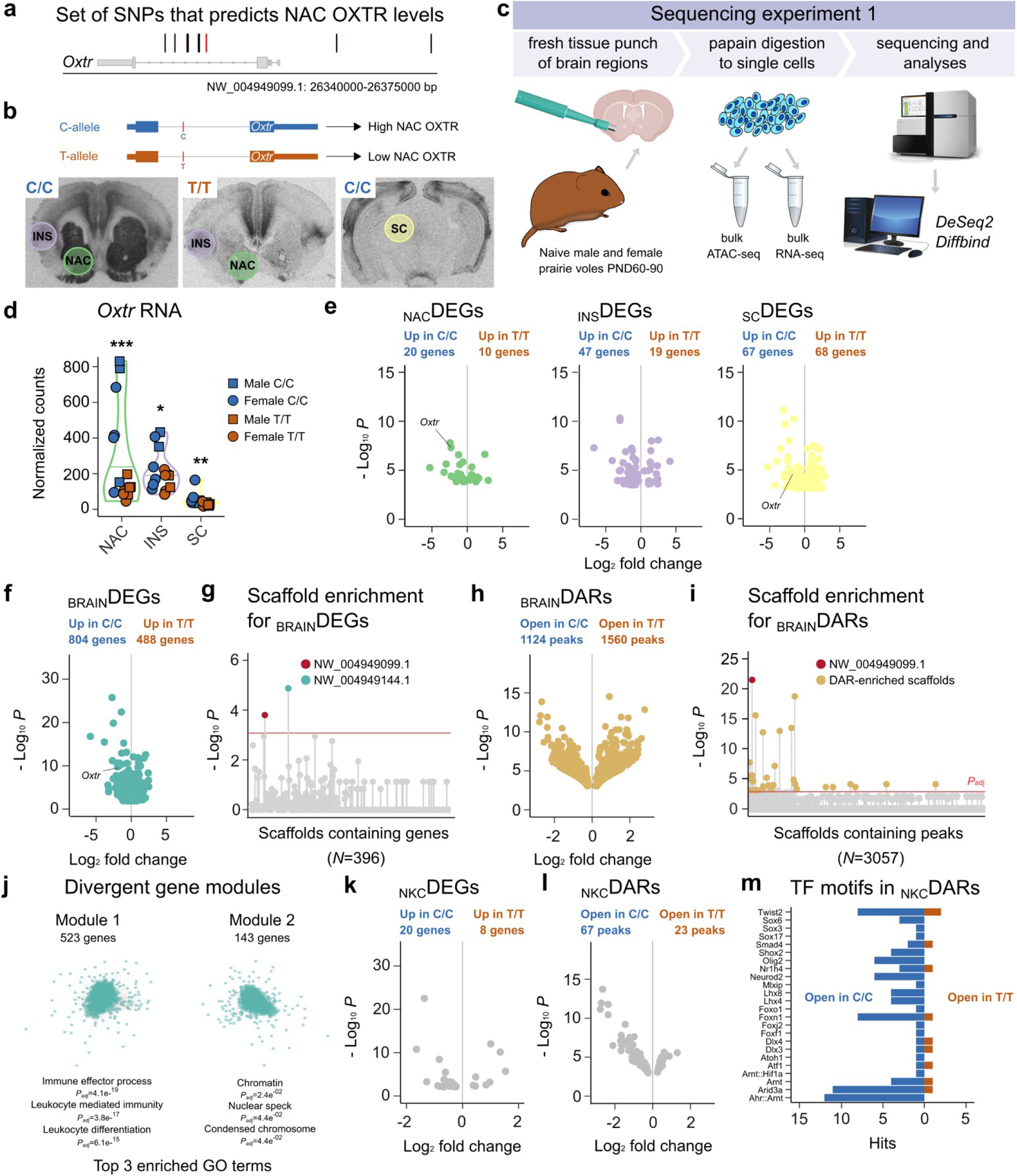
*Oxtr* genotype associates with brain-wide transcriptional changes that are enriched in the natural killer gene complex. **a)** Illustration of the genomic location of previously identified SNPs that predict NAC OXTR levels. In red, the SNP that was genotyped to distinguish high NAC OXTR producers (C/C) from animals with low levels of NAC OXTR (T/T). **b)** Examples of I^125^–OVTA autoradiograms of the two genotypes indicating which brain areas were sampled for sequencing. **c)** Schematic that depicts the workflow of experiment 1. **d)** Normalized *Oxtr* RNA counts for each brain area, split by genotype. * indicates *P*<0.01, ** indicates *P*<0.01, *** indicates *P*<0.001 between genotypes. **e)** Volcano plots depicting _NAC_DEGs, _INS_DEGS and _SC_DEGS genes between C/C and T/T animals. **f)** Volcano plot showing _BRAIN_DEGs between C/C and T/T animals. **g)** Plot depicting enrichment of scaffolds for DEGs. **h)** Volcano plot showing _BRAIN_DARs between C/C and T/T animals. **i)** Plot depicting enrichment of scaffolds for _BRAIN_DARs. **j)** Schematic illustrating two gene modules that are significantly different between genotypes, with top 3 GO terms indicated. **k)** Volcano plot showing _NKC_DEGs between C/C and T/T animals. **l)** Volcano plot showing _NKC_DARs between C/C and T/T animals. **m)** Plot depicting hits of transcription factor motifs in _NKC_DARs. In plot f and i, scaffolds above the red line are significantly enriched (*P*_adj_<0.1). Red dot indicates scaffold containing *Oxtr*.

To investigate whether these region-specific genetic effects extend beyond *Oxtr* expression, we performed bulk ATAC-seq and RNA-seq on the same tissue samples from three brain regions, namely the NAC (high genotypic OXTR variation), the INS (no genotypic OXTR variation), and the superior colliculus (SC, where OXTR density is indistinguishable from background levels) (Figure 1A-C). We first validated that chromatin accessibility and gene expression are functionally linked in our dataset. Indeed, ATAC peaks preferentially located to expressed genes (*P=*2.47e^-323^, Poisson Test, Figure S1A).

Given that non-coding SNPs typically influence gene expression through altered chromatin accessibility, we examined differential accessible regions (DARs) among brain regions, while controlling for genotype, sex and batch (*N*=45). Contrary to our expectations, we found no DARs in the *Oxtr* gene locus, nor did we observe any peaks overlapping with the previously identified SNP loci (Figure S1B). These findings suggests that variation in chromatin accessibility within the *Oxtr* gene locus is unlikely to be involved in driving genotype-dependent, region-specific *Oxtr* expression differences.

We next quantified gene expression differences between genotypes in each of these brain regions, while controlling for batch and sex effects (Tables S1A-C). In contrast to OXTR protein distribution patterns^21^, we found genotype-dependent variation in *Oxtr* RNA levels in the NAC (*N*=15, *P*_adj_=5.33e^-08^), in the INS (*N*=14, *P*=0.03) and in the SC (*N*=16, *P*=2.75e^-05^). However, the difference in the INS did not survive adjustment for multiple comparisons, indicating a blunted effect of genotype on *Oxtr* expression in the INS. In addition, low levels of *Oxtr* RNA in the SC do not translate into detectable protein (Figure 1D-E). Taken together, these results agree with our earlier findings that genotype effects on OXTR density are most pronounced in the NAC^21^.

Interestingly, we identified additional genotype-associated differentially expressed genes (DEGs, defined as *P*_adj_<0.1 and |log₂ fold change| > 0.2) in all three brain regions: 30 _NAC_DEGs, 66 _INS_DEGs, and 135 _SC_DEGs. These results suggest broad transcriptional consequences of *Oxtr* genetic variation that extend beyond the *Oxtr* gene and regions with highly variable OXTR abundance.

Indeed, nine genes showed differential expression in all three brain regions, with two of particular interest: *Mb4* and *Klrb1a*, which both localize to the same scaffold as *Oxtr*^39^. In the fragmented prairie vole gene assembly, scaffolds represent computationally constructed DNA sequences that can be thought of as partial chromosome maps. DEGs located to the *Oxtr* scaffold may indicate a genetic linkage to genotype, as they reside to the same genomic as our characterized SNP set. Since our previous study only surveyed genetic variability within ∼30 kb of the *Oxtr* gene locus^21^, additional SNPs in LD may exist outside this region. Such variants could be responsible for the broader genotype effects on transcription observed here, suggesting that the *Oxtr* gene might be part of a larger genetic module that influences multiple genes simultaneously.

### *Oxtr* genotype associates with clusters of DEGs and DARs suggestive of coordinated functional regulation

To increase our statistical power for identifying genes that consistently vary between genotypes across brain regions (_BRAIN_DEGs), we collapsed data across NAC, INS and SC, while controlling for region, sex, and batch. This combined analysis provided strong evidence for divergent gene expression between genotypes as 1292 _BRAIN_DEGs were identified (*N*=45, _BRAIN_DEGs, Figure 1F and Table S1D). A salient pattern emerged from this analysis: 42 _BRAIN_DEGs located to the *Oxtr* scaffold (NW_004949099.1). To determine whether this clustering was statistically significant, we performed Fisher’s Exact Tests across all genomic scaffolds. This analysis identified 2 scaffolds significantly enriched for DEGs (*P*_adj_<0.1, Fisher’s Exact Test), with the *Oxtr* scaffold showing clear enrichment (*P*_adj_*=*0.03, Fisher’s Exact Test, Figure 1G and Table S1E). This result indicates that a disproportionate number of genes in the genomic vicinity of *Oxtr* are differentially expressed between genotypes, strongly suggesting that genotype effects on transcription are linked to the *Oxtr* scaffold.

We next applied the same approach to our ATAC-seq data to investigate whether any scaffolds were enriched for _BRAIN_DARs (differential accessible between genotypes across brain regions, *P*_adj_<0.1 and |log₂ fold change| > 0.2). *Oxtr* genotype was associated with 2684 differentially accessible regions (Figure 1H and Table S1F). Fisher’s Exact Tests revealed that 36 scaffolds were significantly enriched for _BRAIN_DARs, with the *Oxtr-*containing scaffold demonstrating the strongest enrichment (*P*_adj_*=*9.4e^-19^, Figure 1I and Table S1H).

Together, these complementary analyses reveal strong differences between genotypes in both gene expression and chromatin accessibility in the relative vicinity of the *Oxtr* gene locus. The parallel enrichment patterns suggest that genotype differences include coordinated regulation of gene clusters on this scaffold. However, the mechanistic basis for how these distinct genotypes drive such widespread differences in gene expression remains to be determined.

### The natural killer gene complex shows strong transcriptional divergence between genotypes

To investigate which biological processes might be differentially affected by genotype, we performed weighted gene correlation network analysis (WGCNA) that identified two divergent gene modules between genotypes (*P*<0.05) (Figure 1J). One module was enriched for gene ontology (GO) terms related to chromosome organization, while the other was linked to immune processes (Tables S1I-L).

Interestingly, the immune module contained *Klrb1a*, which is the top _BRAIN_DEG expressed from the *Oxtr* scaffold (Table S1D). *Klrb1a* encodes a C-Type Lectin-Like Receptor (CTLR)^40^, which are expressed from the NKC. This is a genetic region that regulates innate immunity^41,42^, and locates to the same chromosome as *Oxtr* in rats and mice^43^. Given the location of *Klrb1a* on the outer edge of NW_004949099.1 and the fragmented nature of the prairie vole genome assembly^39^, we next searched for additional scaffolds in the prairie vole genome that contain parts of the NKC sequence.

To identify NKC scaffolds likely to map to the same chromosome as *NW_004949099.1*, we employed a computational approach. We calculated the percentage of annotated genes on each prairie vole scaffold that are present on mouse chromosome 6, which harbors the *Oxtr* gene (Figure S2A). Additionally, we determined which prairie vole scaffolds contain CTLRs (Figure S2B). By mapping these combined results to the mouse NKC^34^, we concluded that part of the OXTR scaffold NW_004949099.1:36120000-41749222 as well as scaffolds NW_004949144.1 and NW_004949240.1 contain parts of the prairie vole NKC region, with NW_004949097.1 flanking the NKC region (Figure S2C).

Next, Fisher’s Exact Tests confirmed that our constructed NKC region is enriched for both _BRAIN_DEGs (*P=*9.33e^-06^) and _BRAIN_DARs (*P=*3.36e^-39^). These results strongly suggest that NKC transcription is linked to genetic variation in the *Oxtr* gene locus. They also align with the results of our WGCNA analysis that linked genotype differences to altered immune processes and chromosome organization. Specifically, we identified 28 DEGs in the NKC region (_NKC_DEGs, 20 up in C/C, 8 up in T/T, Figure 1K), and 90 _NKC_DARs (67 more accessible in C/C, 23 more accessible in T/T, Figure 1L). Consistent with the known immune function of the NKC, we found that binding motifs of transcription factors that are involved in immune signaling, such as AHR:ARNT, ARID3A and FOXN1, were significantly more present in _NKC_DARs open in C/C animals (Poisson Test, *P_adj_*<0.1, Table S1M, Figure 1M)^44–46^. This finding provides further support for divergent immune gene regulation between *Oxtr* genotypes.

### Oxtr genotype associates with gene expression across diverse neuronal and glial populations

While our bulk RNA-seq analyses provide strong evidence for genotype-associated transcriptional differences, they could not distinguish whether these effects were restricted to OXTR-expressing cells or extended to other cell types. To address this question, we used our single nuclei dataset from pooled tissue from the NAC and the paraventricular nucleus (PVN) produced with single nucleus RNA-seq (snRNA-seq, *N*=16, Figure 2A-B). We previously profiled these areas because the NAC has variable OXTR expression between genotypes while the PVN harbors OXT-producing neurons.

**Figure 2.**
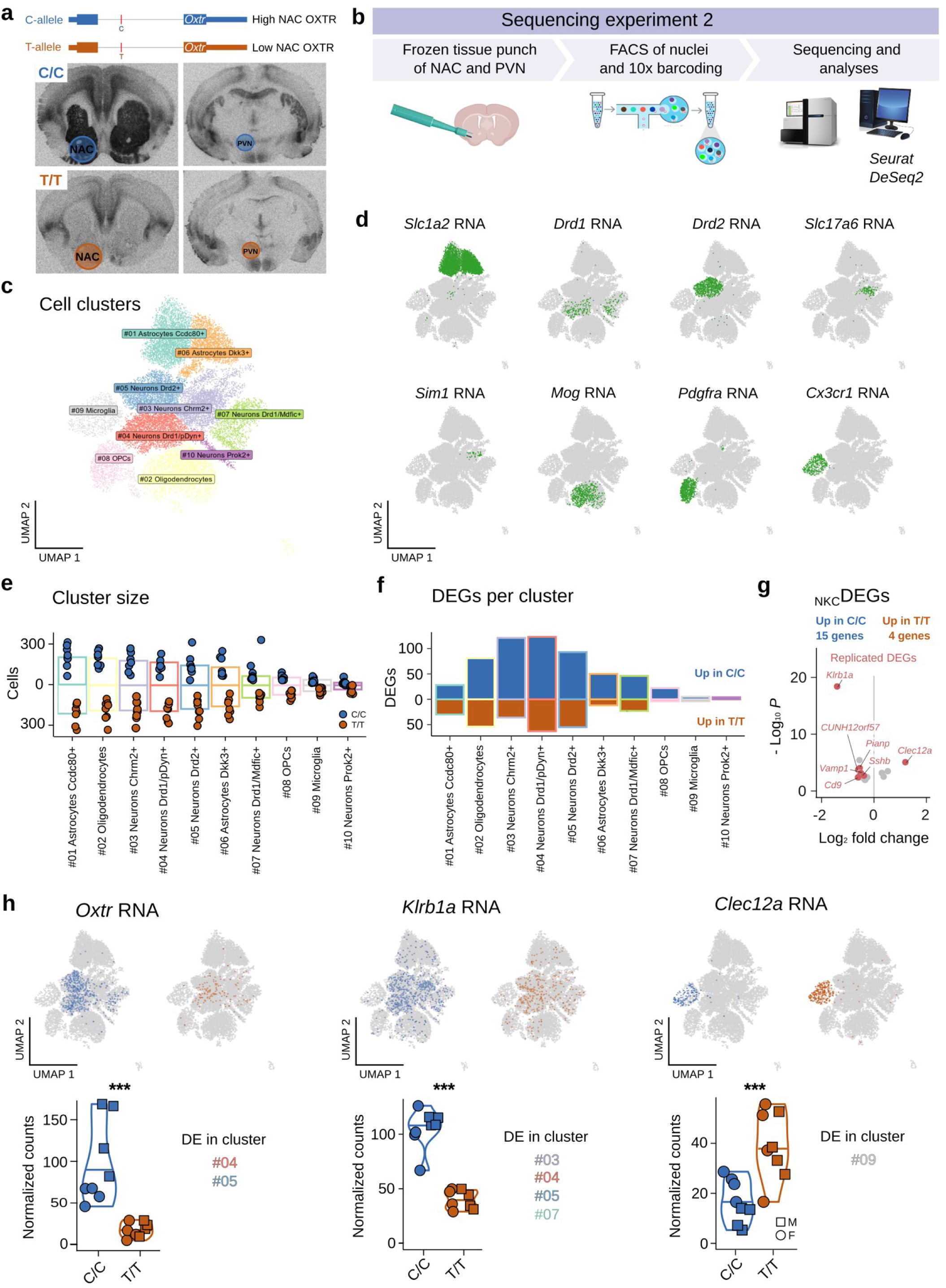
*Oxtr* genotype associates with transcriptional differences across diverse cell types, including altered NKC transcription in OXTR+ neurons and microglia. **a)** Examples of I^125^–OVTA autoradiograms of the two genotypes indicating which brain areas were sampled for sequencing. **b)** Schematic of workflow of experiment 2. **c)** Schematic of annotated cell clusters. **d)** UMAP plots depicting RNA expression of markers used for cluster annotation **e)** Size of cell clusters, split by genotype. **f)** Number of DEGs per cell cluster, split by genotype. OPCs are oligodendrocyte precursor cells **g)** Volcano plot showing _NKC_DEGs between C/C and T/T animals. Red dots represent replicated _NKC_DEGs **h)** Expression patterns and counts of *Oxtr* (*Left*) and the two top _NKC_DEGs: *Klrb1a* (*Middle*) and *Clec12a* (*Right*). UMAP plots depict RNA expression with cellular resolution across animals, violin plots show bulk expression levels per animal. Indicated are the cell clusters in which genes were DE in pseudo-bulk analysis. *** indicates *P*_adj_<0.001 in bulk analysis.

Using the *Seurat* package^47^, we clustered cells into 10 distinct clusters (Figure 2C), which we annotated with established gene markers that strongly differentiated among major cell classes (e.g., neurons and glia) and subclasses (e.g., D1/pDyn+, D1/Mdfic+ and D2+ neurons, Figure 2D and Table S2A). Importantly, cluster size was not significantly different between genotypes (*P*=0.59, RM2W-ANOVA, Figure 2E), indicating that genotype does not alter overall cell-type composition and enabling valid comparisons of gene expression differences between genotypes with cellular resolution.

We quantified transcriptional differences between genotypes using a pseudo-bulk analysis, while controlling for sex. Notably, we found evidence of differential gene expression in all cell clusters (Figure 2F, Table S2B). As expected, *Oxtr* expression differed significantly between genotypes in the *Drd1/pDyn*+ (*P_adj_*=4.97e^-09^) and *Drd2*+ cell clusters (*P_adj_*=7.79e^-20^), consistent. with our previous findings that NAC *Oxtr* mRNA predominantly co-localizes with dopamine receptor mRNAs (*Drd1* and *Drd2*)^48^.

The largest numbers of *Oxtr*-genotype-associated DEGs were observed in cell types that strongly express *Oxtr* (>100 DEGs in both *Drd1/pDyn*+ and *Drd2*+ neurons). However, substantial numbers of DEGs were also observed in oligodendrocytes (>100) and *Dkk*+ astrocytes (>50), cell types in which no to very little *Oxtr* expression could be detected. This finding further reinforces that genotype affects gene expression in a widespread manner, even extending to cell types that do not show genotype-associated variation in *Oxtr* RNA levels, such as glia.

To validate our previous finding that the NKC region is enriched for DEGs, we determined DEGs across cell clusters, as if it were a bulk sequencing experiment (_BRAIN_DEGs, Table S2C). Consistent with our previous bulk RNA-seq results, the NKC region was significantly enriched for _BRAIN_DEGs (*P* = 0.024). In total, 19 out of 1571 _BRAIN_DEGs were _NKC_DEGs (15 up in C/C, 4 up in T/T).

While 7 _NKC_DEGs were replicated from our bulk sequencing experiment, 12 out of 28 identified _NKC_DEGs in experiment were not detected in this data set, likely due to the lower sequencing depth characteristic of snRNA-seq experiments. Interestingly, the two top replicated _NKC_DEGs were CTLRs: *Klrb1a* and *Clec12a.* Given that CLEC and KLR proteins form ligand-receptor pairs that modulate natural killer (NK) cell function in the periphery^49^, their coordinated expression raise the possibility of a functional interaction in the brain.

Consistent with this, we observed striking cell-type specific expression for these two CTLRs (Figure 2H). *Clec12a* showed near-exclusive expression in microglia while *Klrb1a* demonstrated preferential expression in *Oxtr-*expressing cells (*P*<9.24e^-16^, Fisher’s Exact Tests), and this complementary expression pattern is consistent with functional crosstalk between microglia and *Klrb1a/Oxtr+* neurons in the NAC. These results consolidate the link between NKC transcription and genetic variation in the *Oxtr* gene locus and hint upon potential mechanisms through which immune-related genes might influence neural circuits.

While our results demonstrate clear associations between *Oxtr* genotype and NKC gene expression - a critical question remains. Are these effects solely driven by genetic linkage, or are they mediated by functional variation in OXTR signaling? Understanding this distinction is essential for determining whether OXTR signaling mediates genotypic effects on transcription.

### Genetic deletion of OXTR reveals that OXTR signaling mediates genotype effects on NKC transcription

To determine whether variation in OXTR abundance drives NKC transcription, we performed bulk RNA-seq on NAC tissue from three groups: C/C, T/T and systemic OXTR knock-out voles with a C/C genetic background (C^Δ^/C^Δ^, Figure 3A-B)^50^. This experimental design enables us to distinguish between effects mediated by functional OXTR signaling versus those attributable to genetic linkage (Figure 3C).

**Figure 3.**
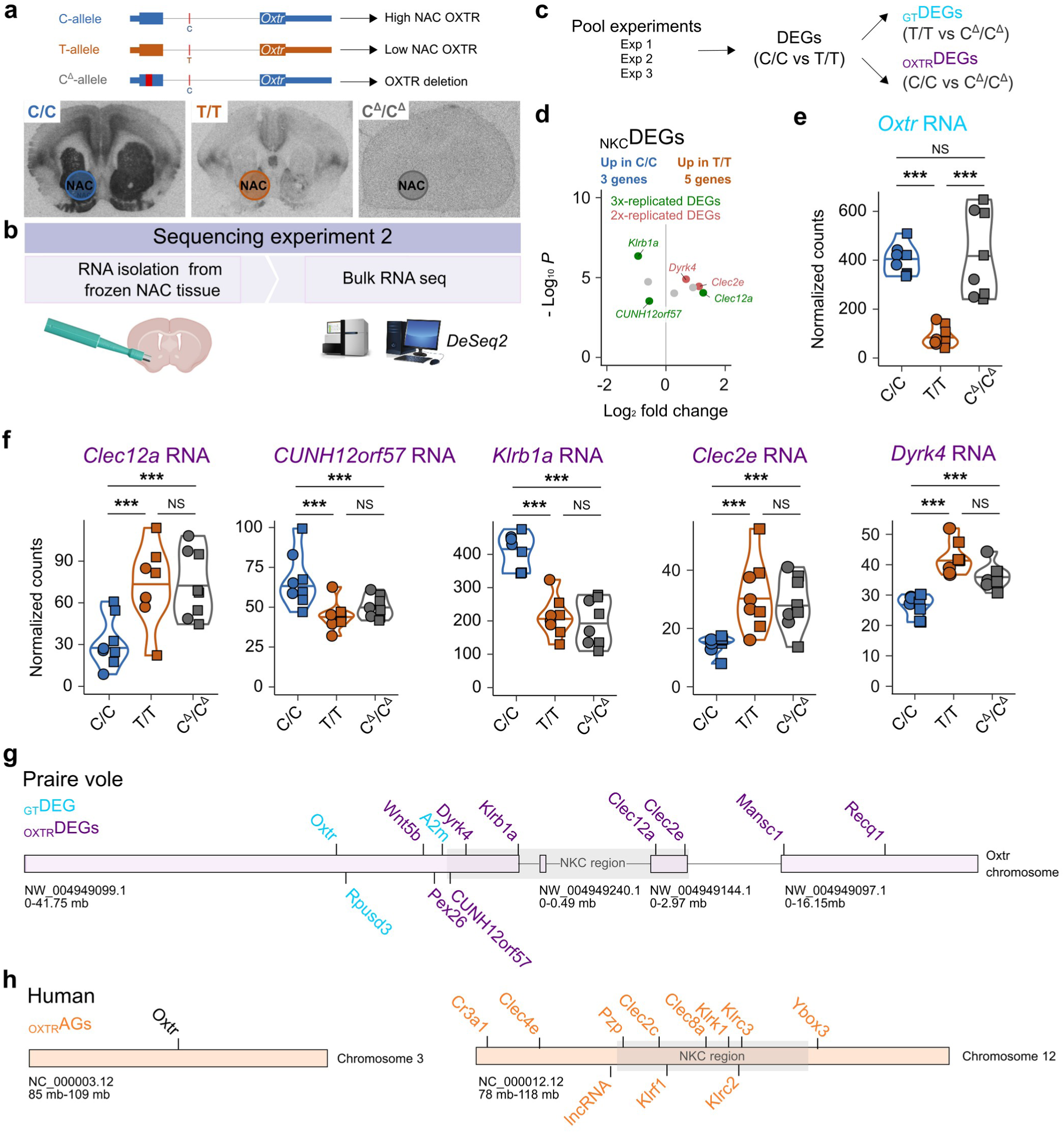
OXTR signaling robustly modulates NKC transcription. **a)** Examples of I^125^–OVTA autoradiograms of the three genotypes indicating the NAC that was sampled for sequencing. **b)** Schematic that depicts the workflow for experiment 3. **c)** Illustration that demonstrates the approach to determine which DEGs were OXTR-mediated versus those solely driven by genotype. **d)** Volcano plot showing _NKC_DEGs between C/C and T/T animals. Red dots represent 2x-replicated _NKC_DEGs, green dots represent 3x-replicated _NKC_DEGs. **e)** Normalized counts of *Oxtr* RNA, *** indicates *P*<0.001 between genotypes. Normalized counts of replicated _NKC_DEGs: *Clec12a, CUNH12orf57*, *Klrb1a*, *Clec2e* and *Dyrk4* RNA, *** indicates *P*<0.001 between genotypes. **f)** Schematic of the constructed prairie vole *Oxtr* scaffold with gene locations. In light-blue are _GT_DEGs, in purple are _OXTR_DEGs. The location of the NKC is shaded gray. **g)** In humans, *Oxtr* and the NKC are not located to the same chromosome. However, the NKC (shaded gray) is enriched for genes that correlate with varying *OXTR* mRNA levels – or _OXTR_AGs, which are depicted in orange.

Initial validation confirmed our previous results: comparing C/C versus T/T animals (*N*=16), while controlling for sex, revealed 76 DEGs, with *Oxtr* showing the strongest association (*P_adj_*=3.84e^-13^) (Table S3A), The NKC region remained significantly enriched for DEGs (*P*=4.55e^-08^, Fisher’s Exact Test, Figure 3D), with 8 _NKC_DEGs identified (3 up in C/C, 5 up in T/T). Notably, 3 _NKC_DEGs replicated across all sequencing datasets (*Clec12a*, *CUNH12orf*57, *Klrb1a*). 2 additional _NKC_DEGs (*Clec2e* and *Dyrk4*) were only detected in the bulk RNA-seq experiments, likely due to the greater sequencing depth of the bulk RNA-seq experiments compared to snRNA-seq.

To distinguish between genotype-driven and effects mediated by OXTR-signaling, we developed a classification system using the C^Δ^/C^Δ^ animals. We designated DEGs as OXTR-mediated (_OXTR_DEGs) when they were differentially expressed in C/C versus C^Δ^/C^Δ^, but not in T/T versus C^Δ^/C^Δ^ comparisons. Conversely, we classified DEGs as genotype-driven but OXTR-independent (_GT_DEGs) when they were differentially expressed between T/T versus C^Δ^/C^Δ^, but not between C/C and C^Δ^/C^Δ^ animals.

Importantly, we confirmed that C^Δ^/C^Δ^ animals harbor a 60 base pair deletion in the first exon of *Oxtr* transcripts (Figure S3), and our previous work demonstrated that this mutation completely abolishes receptor binding and downstream signaling^50^. Therefore, while *Oxtr* is expressed in C^Δ^/C^Δ^ animals, the resulting RNA does not translate into functional OXTR protein, enabling us to determine which genotype effects on transcription require functional OXTR signaling.

The *Oxtr* gene itself exemplified a _GT_DEG: C^Δ^/C^Δ^ animals produced similar levels of (truncated) NAC *Oxtr* RNA as C/C animals, indicating that genotypic effects on *Oxtr* expression are independent of OXTR signaling (Figure 3E). In striking contrast, all 5 replicated _NKC_DEGS were OXTR-mediated (*P<0.05*). Deletion of OXTR significantly affected gene expression levels such that the expression levels of these _NKC_DEG in C^Δ^/C^Δ^ animals mirrored expression levels in T/T voles (Figure 3F, Tables S3B-C). To expand this analysis beyond the NKC, we used Fisher’s method to combine the p-values of genes that showed directionally consistent fold changes (|log₂ fold change| > 0.2) and significance (*P*<0.05) across the three independent C/C vs T/T comparisons. This approach allowed us to include genes with subtle, but consistent brain-wide differences between genotypes. After multiple comparison corrections, we identified 64 high-confidence DEGs (_HC_DEGs) that were consistently and significantly different between C/Cs and T/T animals (Table S3D). Among these _HC_DEGs, we found 20 _OXTR_DEGs and 8 OXTR-independent _GT_DEGs. The _GT_DEGs included 3 genes that localize to the *Oxtr* scaffold, but not to the NKC region: *Oxtr, Rpusd3* and *A2m* (Figure 3G and Table S3E-F). Since *Rpusd3* and *A2m* are unaffected by a loss of OXTR signaling, they may be affected by the same unidentified genetic module as *Oxtr* and/or causal to individual variation in NAC OXTR densities.

Most importantly, our findings demonstrate that replicated _NKC_DEGs are not genetically linked to variation in the *Oxtr* gene, but to functional OXTR signaling. Overall, functional OXTR signaling is a strong mediator of genotype effects on transcription, with 20 _HC_DEGs classified as _OXTR_DEGs, compared to only 8 _GT_DEGs, demonstrating that functional receptor signaling, rather than genetic linkage, is the stronger determinant of transcriptional differences between genotypes.

### Variation in human NAC *OXTR* expression associates with altered NKC transcription

Our findings in the prairie vole demonstrate that the relation between *Oxtr* genotype and NKC transcription depends on functional OXTR signaling. In humans, *OXTR* and the NKC region are located on different chromosomes (chromosome 3 and 12 respectively)^43^. This raised the intriguing possibility to investigate whether the relationship between OXTR levels and NKC transcriptional activity is present in species in which the NKC and the *Oxtr* locus are not structurally linked.

To investigate this hypothesis, we examined whether variation in human *OXTR* expression correlates with NKC gene expression patterns. We analyzed bulk RNA-seq data from human NAC tissue (*N*=246) available through the GTEx database^28^. To identify genes whose expression significantly correlates with *OXTR* abundance, we used *OXTR* Transcript Per Kilobase Million (TPM) values as the independent variable. This analysis revealed 1379 OXTR-associated genes (_OXTR_AGs, *P*_adj_<0.1 and |log₂ fold change| > 0.2, Table S4). Remarkably, _OXTR_AGs showed significant enrichment in the human NKC region (Chr12:9000000:10600000). 7/31 NKC region genes were _OXTR_AGs (*P=*0.0037, Fisher’s Exact Test), representing a disproportionate concentration of OXTR-associated transcripts. These NKC-localized _OXTR_AGs included six *CTLR*s (*CLEC2C, CLEC8A, KLRC2, KLRC3, KLRF1* and *KLRK1*) and *PZP* (Figure 3H), mirroring the coordinated regulation of NKC transcription identified in our prairie vole experiments.

To determine the cellular localization of these human NKC _OXTR_AGs, we leveraged publicly available snRNA-seq data from human NAC tissue^51^. Although this dataset was enriched for neuronal cells, with limited detection of glial cell types, we observed distinct expression patterns that paralleled our prairie vole findings. *CLEC* family members showed predominant expression in microglia and other glial populations, while *KLR* family receptors were primarily expressed in dopaminoceptive neurons (Figure S5). This complementary cellular distribution is consistent with our prairie vole observations and suggest these receptors enable specific cell-cell interactions.

These findings provide intriguing, yet preliminary support for a conserved relationship between OXTR signaling and NKC transcription that extends from prairie voles to humans. The fact that this association persists despite the chromosomal separation of OXTR and NKC genes in humans supports our findings in prairie voles that functional OXTR signaling, rather than genetic linkage to *OXTR*, regulates OXTR-mediated NKC transcription in the brain.

### Genetic reduction in OXTR levels increases spine density in NAC OXTR+ neurons

Our transcriptional analyses in prairie voles revealed that OXTR signaling regulates the transcription of NKC genes. These included 3 C-type lectin-like receptors that are emerging as key regulators of neural connectivity through regulating microglia-neuron interactions^38,52^, *CUNH12orf57* that encodes protein C10, a regulator of neuronal excitability^37^, and *Dyrk4,* which controls neuronal morphology^53^. Given that genetic deletion of OXTR increases dendritic spine density in mice^54,55^, we hypothesized that these phenomena are mechanistically linked (Figure 4A). If OXTR signaling coordinates NKC gene expression to shape synaptic structures, then genetic reduction in OXTR levels should alter the synaptic architecture of OXTR-expressing NAC neurons.

**Figure 4.**
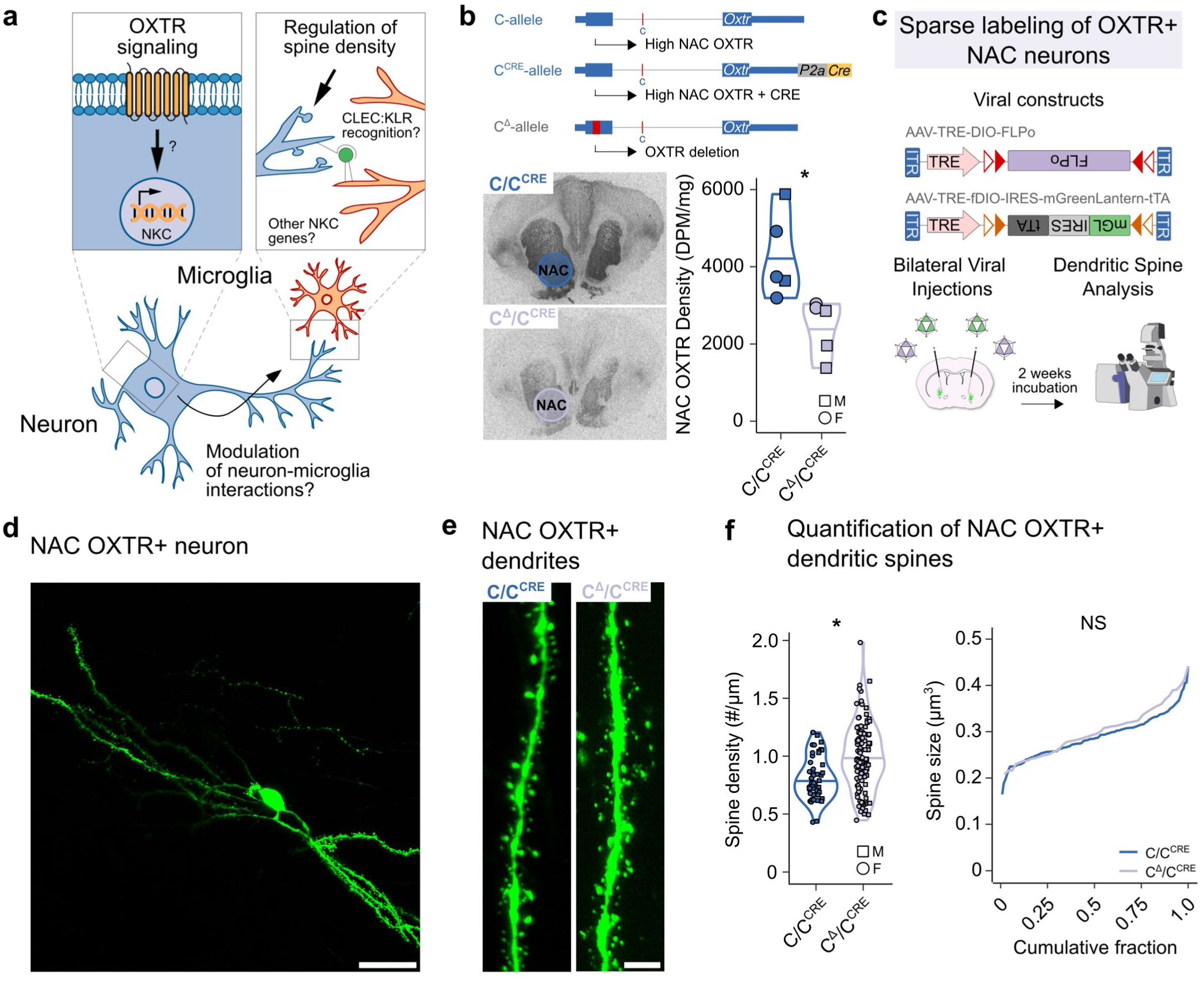
Heterozygous OXTR knockout affects spine density of OXTR+ neurons. **a)** Schematic of how oxytocinergic signaling affects NKC transcription and may regulate synaptic density via modulation of microglia-neuron interactions **b)** Representative autoradiograms and quantification of OXTR densities of the newly created C/C^CRE^ and C /C^CRE^ lines. * indicates *P*<0.05. **c)** Schematic that depicts the experimental workflow. **d)** Representative 2-photon image of sparsely labeled NAC OXTR+ neuron. Scale bar represents 50 μm **e)** Representative 2-photon images of dendrites of either genotype. Scale bar represents 5 μm **f)** Quantification of dendritic spine density and size between genotypes. * indicates *P*<0.05.

To test this hypothesis, while controlling for potential confounding effects of genotype, we employed our recently created OXTR-CRE voles, which selectively express CRE recombinase from the C-allele in OXTR+ cells^56^. We crossed OXTR-CRE animals with OXTR KO prairie voles to generate two experimental groups: C/C^CRE^ animals (with high OXTR levels) and C^Δ^/C^CRE^ animals (with low OXTR levels) (Figure 4B). Autoradiographic analysis confirmed that C^Δ^/C^CRE^ prairie voles exhibited significantly lower, but still detectable, levels of NAC OXTR binding compared to C/C^CRE^ animals (*P*=0.017, Independent Samples T-Test). This reduction mimics the natural variation between C/C and T/T genotypes, providing an ideal system to isolate the effects of OXTR abundance on synaptic architecture.

To specifically examine synaptic changes in OXTR+ neurons, we infused adeno-associated viruses into the NAC to specifically and sparsely label OXTR+ neurons (Figures 4C and S5), enabling high resolution analysis of dendritic spines (Figures 4D-E). Quantitative analysis by means of a linear mixed model controlling for sex and individual revealed that dendritic spine density was significantly increased in OXTR+ neurons of C^Δ^/C^CRE^ compared to C/C^CRE^ animals (Figure 4F, *F_1,11_*=6.9113, *P*=0.0235). Importantly, this increase in spine number occurred without changes in spine size, as this metric was similar between genotypes (*F_1,11_*=0.896, *P*=0.364). These results demonstrate that variation in OXTR abundance is sufficient to alter synaptic architecture of OXTR-positive neurons. The increased spine density observed in OXTR-positive neurons of animals with reduced OXTR levels is consistent with OXTR signaling refining synaptic connectivity. This may represent a mechanism through which natural genetic variation in OXTR expression shapes neural circuit function and, ultimately, behavioral phenotypes.

## Discussion

Through three independent sequencing experiments, we have uncovered that natural variation in OXTR levels links to widespread transcriptional changes that extend beyond the receptor itself. Our findings reveal that genetic variation in the prairie vole *Oxtr* gene locus affects gene expression even in regions and cell types that do not show genotype-dependent differences in OXTR levels. Some transcriptional effects of genotype occur independently of OXTR signaling, which indicates that these genes are affected by the same unidentified genetic module as *Oxtr* and/or causal to individual variation in NAC OXTR densities. However, our most striking finding is that variation in functional OXTR signaling robustly modulates NKC transcription in prairie voles, with additional preliminary support for a similar relationship in humans.

The NKC region represents one of the most conserved immune gene clusters across mammalian species, traditionally known for its role in enabling natural killer (NK) cells to discriminate self from non-self through CTLRs^35,42^. These receptors recognize diverse protein ligands and orchestrate immune responses by regulating cytotoxic defense mechanisms against pathogens^57^. However, the NKC contains other immune genes as well, including cytokines, proteases and tyrosine kinases, among others, and the evolutionary conservation of NKC genetic organization suggests coordinated regulation of the immune genes. Our transcriptional analyses consistently identified NKC enrichment for DEGs, with parallel enrichment for DARs. Critically, our knockout experiments demonstrated that all replicated _NKC_DEGs are regulated by functional OXTR signaling rather than genetic linkage, establishing OXTR as a potent modulator of NKC gene transcription. These findings provide a mechanistic foundation for the emerging role of OXTR signaling in neuroimmune pathways^58–60^.

The functional parallels between central and peripheral immune recognition systems offer insight into how OXTR signaling may influence neural development. Like peripheral NK cells, microglia use lectin-based recognition systems to interact with their environment^61^, and a microglial CTLR has been shown to play a role in the neural circuit refinement^38^. Our identification of complementary CTLR expression patterns – with *Klrb1a* preferentially expressed in *Oxtr-*expressing dopaminoceptive cells, and *Clec12a* restricted to microglia, suggests that OXTR signaling orchestrates microglia-neuron recognition processes analogous to peripheral immune surveillance^49,57^. Additionally, we identified *CUNH12orf57*, which encodes protein C10 that controls excitatory neuronal homeostasis^37^, and *Dyrk4*, a tyrosine kinase that regulates neuronal morphology^53^. These findings converge with evidence that OXTR signaling influences microglial activity^62^, dendritic spine density (this study), dendritic complexity^54^, and excitatory neurotransmission^63^, suggesting a coordinated pathway through which OXTR modulates neurodevelopmental trajectories by influencing microglia-neuron interactions. Supporting this framework is that social isolation, which reduces OXTR signaling^64^, alters microglial morphology in prairie voles^65^, linking social experience to neuroimmune function. Our demonstration that genetic reduction in OXTR levels increases spine density in OXTR+ neurons provides direct evidence that variation in OXTR density is sufficient to alter dendritic architecture, potentially through NKC-mediated microglia-neuron interactions.

These results represent an important advancement in our understanding how OXTR signaling may affect neural circuit function, by revealing a molecular mechanism via which OXTR signaling exerts anti-inflammatory effects. We know from decades of rodent, primate, and human studies that OXTR signaling has the capacity to modulate emotionality and social behaviors in adult individuals^16^, but the influence of OXTR signaling on neurodevelopment has been much less explored^66^. Our work provides a mechanistic framework for understanding how natural variation in OXTR signaling, in interaction with social environments, shapes the formation of neural circuits underlying social behavior. Previous work in the prairie vole shows that adult social behaviors are susceptible to variation in OXTR signaling interacting with social contexts during critical early life periods^25^. Early-life neonatal separations lead to impairments in social bonding in adult prairie voles with low NAC OXTR, while high NAC OXTR voles are resilient to the same neglect model. Presumably, they are rescued by the heightened OXTR signaling upon reunion with parents afforded by the high OXTR density. Similarly, OXTR genotype and the resulting variation in NAC OXTR density interacts with parental composition (uniparental versus biparental care) to influence adult social attachment^26^. Future work will have to establish whether these gene*environment interactions reflect differential activation of the neuroimmune pathways we have identified here, with increased OXTR signaling in C/C animals providing protection against stress-induced alterations in microglia-neuron interactions.

The translational relevance of these mechanisms is supported by extensive human literature linking genetic variation in the *OXTR* gene to diversity in social and psychiatric outcomes as well as brain functional connectivity. This includes but is not limited to resilience to social neglect^67^, facial recognition^31^, parental behavior^30^, and conditions of altered sociality^29^, such as ASD^33,33,68^. This body of work underscores the impact that individual variation in OXTR signaling, whether mediated by variation in OXTR density or social experience, can have on adult sociality and/or emotionality. However, none of these studies provide mechanistic clues as to how OXTR signaling, in interaction with the social environment, shapes social phenotypes. Our results provide a first step in this direction, by showing that OXTR signaling robustly influences NKC transcription, suggesting a mechanism via which signaling of this nonapeptide affects a neuroimmune pathway to ultimately control neurodevelopmental trajectories and or/responses to stress and the social environment.

In conclusion, our findings demonstrate that natural variation in OXTR signaling modulates neuroimmune pathways and neurodevelopment, providing mechanistic clues into how variation in OXTR signaling acts to shape neural circuits and diverse social phenotypes. These findings open new avenues for further investigating the biological basis of social diversity in health and disease and may ultimately inform approaches to promote social health.

## Methods

### Animal husbandry

All male (*N*=55) and female (*N*=53) prairie voles (*Microtus ochrogaster*) used for experimental were virgins between 60-120 days old (early adulthood) and taken at random from our colony, which consists of breeders derived from field captured voles in Illinois, USA. Prairie voles were housed in same-sex groups (2–3 animals) from post-natal day (PND) 21-23 in ventilated plexiglass cages (36×18×19 cm) filled with bedding and nesting material under a 14:10 h light/dark cycle at 20-22°C and 40-60% humidity with *ad libitum* access to water and food (Lab Rabbit Diet HF #5236, LabDiet). All experimental procedures complied with the ARRIVE guidelines and were approved by the Emory University Institutional Animal Care and Use Committee.

### Genotyping and breeding of C/C, T/T, C^Δ^/C^Δ^, C/C^CRE^ and C^Δ^/C^CRE^

We created breeder pairs producing low- or high nucleus accumbens (NAC) OXTR offspring by randomly genotyping prairie voles from our colony at position NW_004949099.1:26351196. A single-nucleotide polymorphism (SNP; C or T) at this locus strongly predicts NAC *Oxtr* expression. Genomic DNA was isolated from ear clips with the DNAeasy Blood and TIssue kit (Qiagen, Germany). Allele-specific PCR amplification of the SNP locus was achieved using HiDi® DNA Polymerase (myPols Biotec., Germany), allele-specific forward primers (FW_CC: 5’-GTG GAA TCA TCC CAC CGT GC -3’, FW_TT: 5’ - GTG GAA TCA TCC CAC CGT GT -3’) and a reverse primer (RV:5’-TCC CAA TGT GCT AGA GCA GG -3’). Cycling conditions consisted of initial denaturation: 95 °C for 2 min, 35 amplification cycles: 95 °C for 15 sec, 53 °C for 15 sec, 72 °C for 30 sec, and a final elongation phase: 72 °C for 5 min. Heterozygous (CT) animals were selected for further breeding. Their homozygous offspring (either CC or TT) were subsequently crossed to create breeder pairs to solely produce low *Oxtr* (TT)- or high *Oxtr* (CC)-expressing animals. For genotyping the OXTR KO allele (C^Δ^), we PCR-amplified the region around the 60 bp deletion in the *Oxtr* gene using these primers (FW: AGATCAGTGCCCGGGTGCCC, RV: TCGAGCGACATAAGCAGCAG). Cycling conditions were: 95 °C for 2 min, 35 amplification cycles: 95 °C for 15 sec, 53 °C for 15 sec, 72 °C for 30 sec, and a final elongation phase: 72 °C for 5 min. Heterozygous OXTR KO animals were first bred with either CC or TT animals. Subsequently, their heterozygous OXTR KO offspring were crossed (C/C^Δ^xC/C^Δ^ and T/C^Δ^xT/C^Δ^ breeder pairs) such that their combined offspring would include TT, CC and C^Δ^/C^Δ^ animals. For genotyping the OXTR CRE allele (C^CRE^), we PCR-amplified the region around the 60 bp deletion in the *Oxtr* gene using these primers (FW: TGCCTTCATCATCGCCATGC, RV: CGACAGGCGCGGTGGCCAGA, RV: CCGCGCTGGAGTTTCAATAC). Cycling conditions were: 95 °C for 2 min, 35 amplification cycles: 95 °C for 15 sec, 53 °C for 15 sec, 72 °C for 30 sec, and a final elongation phase: 72 °C for 5 min. Heterozygous OXTR KO animals (C/C^Δ^) were bred with C^CRE^/C^CRE^ animals, such that their offspring was C/C^CRE^ or C^Δ^/C^CRE^. Care was taken to prevent inbreeding; all mates had a kinship coefficient of <12.5%

### I^125^–OVTA autoradiography

Fresh frozen brains were sectioned on a cryostat at 20 µM and mounted on Fisher Superfrost plus slides and stored at -80 °C. The slides were thawed and fixed for two min with 0.1% paraformaldehyde in PBS at RT and then incubated in 50 pM I^125^–OVTA (2200 Ci/mmol; PerkinElmer; Boston, MA), a selective, radiolabeled OXTR ligand, for one hour. Unbound I^125^–OVTA was then removed with Tris-MgCl_2_ buffer and sections were allowed to dry. Sections were exposed to BioMax MR film for 7 days, after which films were developed, and images were taken using an MCID camera set-up (InterFocus Ltd, UK). For quantification, mean gray values, corrected for background, were determined in ImageJ. I^125^–activity (disintegrations per minute) was calculated using an I^125^–standard and taken as a proxy for OXTR density. Differences in OXTR density were determined by comparing NAC OXTR protein density levels of C/C^CRE^ and C^Δ^/C^CRE^ animals.

### Bulk total RNA and ATAC sequencing sample preparation

Animals were sacrificed using CO_2_ asphyxiation, after which brains were dissected out and sliced on a ice-cold mouse brain matrix (Zivic Instruments, USA). For combined RNA and ATAC sequencing, slices were then transferred to ice-cold HBSS and nucleus accumbens (NAC), insular cortex (INS) and superior colliculus (SC) were isolated using micro punches (1.2 and 2 mm diameter). Punches were digested with papain dissociation enzyme (Thermo-Scientific, USA) in 250 ul HBSS containing 500 units of DNAse (Sigma-Aldrich, Germany) at 37 °C for 30 minutes. Cells were mechanically dissociated and filtrated into single-cell solution using FlowMi filters (40 and 70 um, BelArt, USA). Viability and yield were checked by tryptophan staining after which cells were cryopreserved in HBSS supplemented with 10% DMSO and stored at -80 °C before sending for sequencing (Genewiz, LLC., NJ, USA). Each sample was both ATAC-and RNA-sequenced, except for samples CMI and TMI, that were only ATAC-sequenced. For bulk RNA sequencing of the OXTR KO animals, NAC punches were made from flash-frozen brain slides, stored at -80 °C and sent for sequencing (Genewiz. LLC).

### Bulk total RNA sequencing library preparation and processing

Total RNA was extracted from samples using a standard TRIzol extraction method. RNA samples were quantified using Qubit 2.0 Fluorometer (Life Technologies, Carlsbad, CA, USA) and RNA integrity was checked using Agilent TapeStation 4200 (Agilent Technologies, Palo Alto, CA, USA). rRNA depletion sequencing library was prepared by using QIAGEN FastSelect rRNA HMRKit (Qiagen, Hilden, Germany), per the manufacturer’s instructions. RNA sequencing library preparation uses NEB Next Ultra II RNA Library Preparation Kit for Illumina by following the manufacturer’s recommendations (NEB, Ipswich, MA, USA). Briefly, enriched RNAs are fragmented for 15 minutes at 94 °C. First strand and second strand cDNA are subsequently synthesized. cDNA fragments are end repaired and adenylated at 3’ends, and universal adapters are ligated to cDNA fragments, followed by index addition and library enrichment with limited cycle PCR. Sequencing libraries were validated using the Agilent Tapestation 4200 (Agilent Technologies, CA, USA), and quantified using Qubit 2.0 Fluorometer (ThermoFisher Scientific, MA, USA) as well as by quantitative PCR (KAPA Biosystems, MA, USA). The sequencing libraries were clustered on 1 lane of a flowcell. After clustering, the flowcell was loaded on an Illumina HiSeq instrument, according to manufacturer’s instructions. The samples were sequenced using a 2×150bp Paired End (PE) configuration. Image analysis and base calling were conducted by the HiSeq Control Software (HCS). Raw sequence data (.bcl files) generated from Illumina HiSeq was converted into fastq files and de-multiplexed using Illumina’s bcl2fastq 2.17 software. One mismatch was allowed for index sequence identification.

### Bulk total RNA-seq processing and DEG identification

Read quality was checked with *FastQC* and adapter sequences were trimmed with Cutadapt^69^. Reads were aligned to the prairie vole genome (GCA_000317375.1) with *STAR 2.7.10a* in default mode^70^. *FeatureCounts* was used count reads within genes (exonic and intronic), and genes that had >5 counts in >22 samples were input to *Deseq2* to identify differentially expressed genes (DEGs)^71,72^.The model used to look at region-specific effects of genotype was ∼group+sex+batch, where group had six levels (NAC_CC, NAC_TT, INS_CC, INS_TT, SC_CC and SC_TT), while for differential expression analysis between genotypes the model ∼genotype+area+sex+batch was used. For the analysis of KO animals, genes with >5 counts in >11 individuals were included and the model ∼genotype+sex was used. Genes were differentially expressed if *P_ad j_*<0.1 and l |log₂ fold change| > 0.2.

### ATAC sequencing library preparation and processing

Cryopreserved cell samples were thawed, washed, and treated with DNAse I (Life Tech, CA, USA) to remove genomic DNA contamination. Cell samples were quantified and assessed for viability using a Countess Automated Cell Counter (ThermoFisher Scientific, Waltham, MA, USA). After cell lysis and cytosol removal, nuclei were treated with Tn5 enzyme (Illumina, CA, USA) for 30 minutes at 37°C and purified with Minelute PCR Purification Kit (Qiagen) to produce tagmented DNA samples. Tagmented DNA was barcoded with Nextera Index Kit v2 (Illumina) and amplified via PCR prior to a SPRI Bead cleanup to yield purified DNA libraries. The sequencing libraries were clustered on a single lane of a flowcell. After clustering, the flowcell was loaded on the Illumina HiSeq according to manufacturer’s instructions. The samples were sequenced using a 2×150bp Paired End (PE) configuration. Image analysis and base calling were conducted by the HiSeq Control Software (HCS). Raw sequence data (.bcl files) generated from Illumina HiSeq was converted into fastq files and de-multiplexed using Illumina’s bcl2fastq 2.17 software. One mismatch was allowed for index sequence identification.

### ATAC sequencing data processing, DAR identification and transcription factor motif analysis

Read quality was checked with *FastQC* and adapter sequences were trimmed with *Cutadapt*^69^. Reads were aligned to the prairie vole genome (GCA_000317375.1) with *Bowtie2* with the *--very-sensitive* option enabled^73^. Next, regions of open chromatin were determined with *Genrich* using *-j -y -r -v* options and a cut off value of *P* < 0.05. Peak sets were constructed by merging samples from each genotype with a significance cutoff *P*_adj_=0.1. These peak sets served as input to *Diffbind*, which was used to calculate differences in chromatin accessibility between brain areas and genotypes, using the model -genotype+area+sex+batch^74^. Regions were considered differentially accessible if *P*_adj_<0.1. For enrichment of transcription factor motifs we extracted the sequences of differential accessible regions and used the package *TFBSTools* in combination with the *JASPAR2022* mouse transcription factor database to search for enrichment of transcription factor motifs with minimal score set at 99%.

### Weighted gene correlation network analysis

Modules of co-expressed DEGs were analyzed using weighted gene correlation network analysis (WGCNA) using the “WGCNA” package in R^75^. The normalized gene expression data for DEGs was used as the input gene expression matrix and the function *pickSoftThreshold* was used to determine the optimal soft-thresholding power. We determined the optimal soft-thresholding power to be 10 because it was the lowest power for which the scale-free topology fit index reached 0.8. Then an adjacency matrix was created from the input gene expression matrix using the adjacency function with power = 10 and otherwise default parameters. The adjacency matrix was transformed into a topological overlap matrix, and modules were detected by hierarchical average linkage clustering analysis for the gene dendrogram. After the modules were identified, the module eigengene (ME) was summarized by the first principal component of the module expression levels. Module–trait relationships were estimated using the correlation between MEs and genotype. To evaluate the correlation strength, we calculated the module significance (MS), which was defined as the average absolute gene significance (GS) of all the genes involved in the module. Modules were considered significantly associated with genotype if *P*<0.05 and visualized with the *iGraph* package. Additionally, we extracted the corresponding gene information for significant modules which was used as in input to *ShinyGO*^76^. For this, we used the genes included in *DEseq2* analysis as background, and limited GO enrichment to pathways that contain between 50 and 500 genes.

### Single nuclei isolation

Animals were sacrificed using CO_2_ asphyxiation, after which brains were dissected out and sliced on a ice-cold mouse brain matrix (Zivic Instruments). NAC and paraventricular hypothalamus (PVN) punches were taken from flash-frozen brain slides and stored at -80 °C. Immediately prior to nuclei isolation, frozen tissue samples were pooled by *Oxtr* genotype and sex, so PVN and NAC tissue were combined. Pooled samples were then incubated in 150 μl chilled lysis buffer containing 10 mM Tris-HCL (Sigma), 10 mM NACl (Sigma), 3 mM MgCl_2_ (Sigma), 0.1% Nonidet P40 Substitute (Sigma), and Nuclease-free H_2_O for 30 minutes with gentle rotation. Following lysis, 150 μl Hibernate AB Complete nutrient medium (HAB; with 2% B27 and 0.5 mM Glutamax; BrainBits), and samples were rapidly triturated using silanized glass Pasteur pipettes (500 μm internal diameter, BrainBits) until complete tissue dissociation. Samples were then centrifuged (400 x g, 5 minutes, 4°C) and resuspended in 1-1.5 ml chilled wash and resuspension buffer containing 2% Bovine Serum Albumin (Sigma) and 0.2 U/μl RNase Inhibitor (Sigma) in 1X PBS (Thermo Fisher). The nuclei suspensions were filtered through 40 μm Flowmi^®^ cell strainers (Sigma) and 30 μm MACS^®^ SmartStrainers (Milltenyi) to remove large debris and aggregations of nuclei prior to fluorescence activated cell sorting (FACS). Singlet nuclei were then further purified using FACS (BD FACSAria^TM^ Fusion Cell Sorter, BD Biosciences) and FACSDiva software (v8.0.1, BD Biosciences). Sizing beads (6 μm; BD Biosciences) and 1.0 μg/ml DAPI (Sigma) were used to set gating parameters, enabling selection of singlet nuclei based on size (forward scatter), shape (side scatter), and DNA content (DAPI fluorescence). Approximately 250,000 nuclei/pool (ranging from 242,421-252,159) were collected into 1 mL wash and resuspension buffer for downstream library preparation and sequencing.

### Single nucleus RNA library preparation and sequencing

Suspensions of purified singlet nuclei were loaded onto the 10x Genomics Chromium Controller (10x Genomics) at concentrations ranging from 240-250 nuclei/ul with a target range of 4,000 nuclei per sample. Downstream cDNA synthesis and library preparation using Single Cell 3’ GEM, Library and Gel Bead Kit v3.1 and Dual Index Kit TT, Set A (PN-1000215/PN-3000431) were performed according to manufacturer instructions (Chromium Single Cell 3’ Reagent Kits User Guide v3.1 Chemistry, 10X Genomics). Sample quality was assessed using high sensitivity DNA analysis on the Bioanalyzer 2100 system (Agilent) and libraries were quantified using a Qubit 2.0 (Invitrogen). Barcoded cDNA libraries were pooled and sequenced on the NovaSeq 6000 platform (Illumina) on a single flow cell (2×150 bp paired end reads; Illumina).

### Single nucleus RNA pre-processing and quality control

FASTQ files were processed with *Cell Ranger* version 3.1.0 (10X Genomics). Reads were aligned to the *Microtus ochrogaster* genome assembly using a splice-aware alignment algorithm (STAR) within *Cell Ranger*, and gene annotations were obtained from the same assembly (NCBI RefSeq assembly accession: GCA_000317375.1, MicOch1.0). Because nuclear RNA contains intronic sequence, they were included in the *Cell Ranger* count step*. Cell Ranger* filtered out UMIs that were homopolymers, contained N, or contained any base with a quality score less than 10. Following these steps, *Cell Ranger* generated four filtered feature-barcode matrices (one per pool) containing expression data for a total of 26,397 features (corresponding to annotated genes) and a total of 21,838 barcodes (corresponding to droplets and putative nuclei) that were used passed through additional quality control steps in the *Seurat* package in R^47^. Total transcripts and total genes varied slightly across pools, and therefore slightly different criteria were used to remove potentially dead or dying nuclei from each pool. Mitochondrial genes were represented at low levels in every pool and were not used as a filtering criterion. To reduce risk of analyzing dead or dying nuclei, barcodes associated with fewer than 400-500 (depending on pool) total genes were excluded from downstream analysis. To reduce risk of doublets or multiplets, barcodes associated with more than 4,000-5,000 total genes and 10,400-18,000 total transcripts were also excluded. This step filtered out a total of 819 barcodes (3.7%).

### Prediction of nuclei from distinct individual animals

To deconvolute nuclei derived from distinct individuals within pools, we used *Souporcell*^77^, a tool that clusters nuclei based on genetic variants detected in RNA reads. Reads were first filtered for base quality scores greater than 10, high confidence mapping to the genome (-F 3844), and exceeding *Cell Ranger* criteria for UMI counting (tag xf:i:25). Next, high-quality reads were used to call variants in each nucleus using the GATK v4.1.8.1 *HaplotypeCaller* function with default parameters, but without the *MappingQualityAvailableReadFilter* to retain reads that were confidently mapped to the genome by *Cell Ranger* (MAPQ score of 255). *Vartrix* v1.1.22 was then used to count alleles in each nucleus (using recommended parameters). Lastly, *Souporcell* v2.0 was used to cluster cells by genotype, using default parameters but with the number of clusters set to 4 based on the known number of individuals in each pool. Thus, four clusters were identified for each pool, with each cluster comprising nuclei that were predicted to have originated from the same individual animal. *Souporcell* also uses clustering information to predict multiplets (based on individual barcodes that are associated with reads assigned to multiple individuals). We filtered out an additional 435 (2%) barcodes that were predicted to be multiplets by *Souporcell*.

### Dimensionality reduction and clustering

To normalize the data, we used the *SCTransform* function in *Seurat*. To examine dimensionality, we first performed a linear dimensional reduction using the *RunPCA* command with the maximum possible number of dimensions (“dim” set to 50). All 50 PCs were used as input for non-linear dimensional reduction (Uniform Manifold Approximation and Projection, UMAP) using the *RunUMAP* function in *Seurat*. For *RunUMAP*, “min.dist” was set to 0.5, “n.neighbors” was set to 50, “spread” was set to 0.2, “n.epochs” was set to 1000, and “metric” was set to “euclidean”. Prior to clustering, nuclei were embedded into a K nearest-neighbor (KNN) graph based on euclidean distance in UMAP space, with edge weights based on local Jaccard similarity, using the *FindNeighbors* function in *Seurat* (reduction=”umap”,k.param=50, dims=1:2, n.trees=500, prune.SNN=0). Global clustering was then performed using the *Seurat FindClusters* function using the Louvain algorithm with multilevel refinement (algorithm=2) with resolution set to 0.2. Clusters that were not present in all individuals (*N*=1) were excluded from further analysis.

### Cluster marker gene analysis

Cluster identities were assigned using a combination of unbiased analysis of cluster-specific marker genes as well as a supervised examination of previously established marker genes. Cluster-specific marker genes were identified using the *FindAllMarkers* function in *Seurat*. Briefly, this function compares gene expression within each cluster to gene expression across all other clusters and calculates Bonferroni-adjusted *p* values using a Wilcoxon rank-sum test. Clusters were annotated based on well-known marker genes that differentiate between major cell-types (neurons, astrocytes, oligodendrocytes, oligodendrocyte precursor cells and microglia) and further subdivided using genes that further separated cell clusters (*e.g.,* D1-neurons and D2-neurons).

### Identification of DEGs in single nucleus RNA sequencing experiment

We used a pseudo-bulk analysis based on *DeSeq2* to identify differentially expressed genes (DEGs) between genotypes in each cluster. Genes that had >5 counts in >8 individuals were included in the analysis. For each independent cell cluster, the model ∼genotype+sex was used, and P-values were adjusted for multiple comparisons across cell clusters. For comparison with bulk RNA sequencing experiments, a standard *DeSeq* analysis was performed across all cells, not taking cluster into account, with the model ∼genotype+sex. Gene were significant if *P*_adj_<0.1 and |log₂ fold change| > 0.2.

### Identification of *OXTR-*correlated genes and *CTLR* expression in human NAC

To identity genes of which the expression correlates with *OXTR* in the human NAC, we used human GTEx bulk RNA sequencing data containing raw gene read counts from NAC tissue, which was aligned to GRCh38/hg38 using *STAR v2.5.3a*, based on the GENCODE v26 annotation. Genes that showed >10 counts in >132 individual were included. *OXTR* TPM values were used as the independent variable in a *DeSeq2* analysis. Genes were considered significantly correlated with *OXTR* if *P*_adj_<0.1 and |log₂ fold change| > 0.2. To determine cell-type specific NAC expression of *OXTR*-correlated genes localized in the NKC we used the publicly available *Seurat* object from Tran *et al.,* with existing cluster annotation^51^. We used the *plotExpression* function to plot raw counts in each NAC cell type.

### In situ hybridization

Fresh-frozen brains (N=2) were sectioned at 20 μm using a cryostat (Cryostar NX-70), mounted on Superfrost Plus slides (Fisher Scientific), and stored at −80°C until use. RNAscope in situ hybridization was performed according to the manufacturer’s protocol (#323100, ACD Inc., MN, USA). Briefly, thawed slides were pretreated with Protease IV (#322380, ACD Inc.) and subsequently incubated with CRE and OXTR probes (#312281, #500721, ACD Inc.) for 2 hours at 40°C. Following washes, signal amplification was performed using the kit’s reagents. Finally, sections were stained with DAPI and coverslipped using ProLong Gold Antifade Mountant (Thermo Fisher Scientific). Images were acquired at 40× magnification using a Keyence BZ-X700 microscope (Keyence, Japan).

### Synthesis of viral vectors

Adeno-associated virus (AAV) particles were synthesized with pAAV-TRE-DIO-FLPo (Addgene, plasmid #118027, gifted by M. Luo) and pAAV-TRE-fDIO-IRES-eGFP-tTA (Addgene, plasmid #118026, gifted by M. Luo), of which we swapped the eGFP sequence for mGreenLantern (mGL), which is a reporter specifically designed for spine quantification^78^. Viral particles were produced in human embryonic kidney (HEK) 293T cells, purified with AVB-affinity chromatography (HiTrap^TM^ AVB Sepharose^TM^ HP Columns, Cytiva, MA, USA) and concentrated by centrifugal filtration (Amicon Ultra-4, Fisher Scientific, NH, USA), after which viral titer was determined using quantitative PCR targeting the inverted terminal repeats. Two viruses were generated: AAV9-TRE-DIO-FLPo and AAV9-TRE-fDIO-IRES-mGL-tTA. These vectors were diluted to 1.0×10^9^ and 2.5×10^7^ genomic copies/μl respectively and mixed in a 1:1 ratio.

### Intracranial surgeries and dendritic spine analysis

Anesthesia was induced by exposure to a 2-4% isoflurane/oxygen mix and maintained at 1-3%. Three daily doses of meloxicam (2 mg/kg) were administered after surgery. Using a stereotaxic apparatus, animals were bilaterally injected with the mix of AAV9-TRE-DIO-FLPo and AAV9-TRE-fDIO-IRES-mGL-tTA. Stereotaxic coordinates were as follows: AP +1.7, ML ±2.1, DV 4.7. Two weeks after surgery, Animals were deeply anesthetized and transcardially perfused with PBS, followed by PBS supplemented with 4% paraformaldehyde (PFA). Brains were dissected out, post-fixed in 4% PFA for 24 hours and stored in PBS at 4°C. Brains for specificity of CRE-expression were sectioned in a sliding microtome at 50 uM, coverslipped in Fluoromount-G containing DAPI and 10X images were taken on a Keyence BZ-X700 microscope (Keyence). Brains for spine analysis were sectioned along the coronal plane at 150 µm using a vibrating microtome (Leica VT1200S). Sections were mounted on glass slides in Fluoromount-G medium (Thermo Fisher Scientific). Dendritic spine analysis was subsequently performed as previously described^79,80^. Briefly, imaging of dendritic spines was performed on a Leica Stellaris confocal microscope using an HC PL APO CS2 63x/1.40 Oil objective (Leica, Germany). Next, image stacks were up-sampled in FIJI by a factor of 2 and automated analysis of dendritic spines was performed using the RESPAN v0.9.91 pipeline (https://github.com/lahammond/respan), a graphical interface-based tool that integrates deep learning for neuron and spine segmentation^80^. A customized nnU-Net model was trained within the RESPAN environment using representative fluorescence image volumes collected from related experiments. The model was then applied through the RESPAN GUI to segment dendritic branches, spine necks, and spine heads in 3D. RESPAN’s segmentation output was used to extract quantitative morphological features including spine density and spine head volumes.

## Acknowledgements

We thank Lorra Julian and the Emory Primate Center veterinary staff for their dedicated animal care. We thank Zakia Sathi for help with genotyping. We also commemorate our dear mentor Larry Young, who sadly passed away during the preparation of this manuscript. We vividly remember his relentless enthusiasm for all things oxytocin, and we are very proud to contribute to his extensive legacy with this work.

## Author contributions

Conceptualization: AJB and LJY. Contribution of KO and CRE animals: KH. Wet-lab experiments for bulk ATAC and RNA-seq: AJB. Wet-lab experiments for snRNA-seq: AJB, BEH and ZVJ. RNA-scope: KI. Analyses of bulk ATAC and RNA-seq: AJB and HW. Analyses of snRNA-seq: AJB, GWG and ZVJ. Design, synthesis and infusion of viral vectors: AJB. Dendritic spine analysis: KKG and ERES. Supervision: AJB, ERES, JTS and LJY. Writing - draft: AJB and LJY. Writing - review and editing: all authors.

## Declaration of generative AI in scientific writing

During the preparation of this work the authors used Claude (Anthropic, USA) and ChatGPT (OpenAI, USA) to improve grammar and readability. After using these tools, the author reviewed and edited the content as needed and take full responsibility for the content of the publication.

## Supplementary figures

**Supplementary figure 1.**
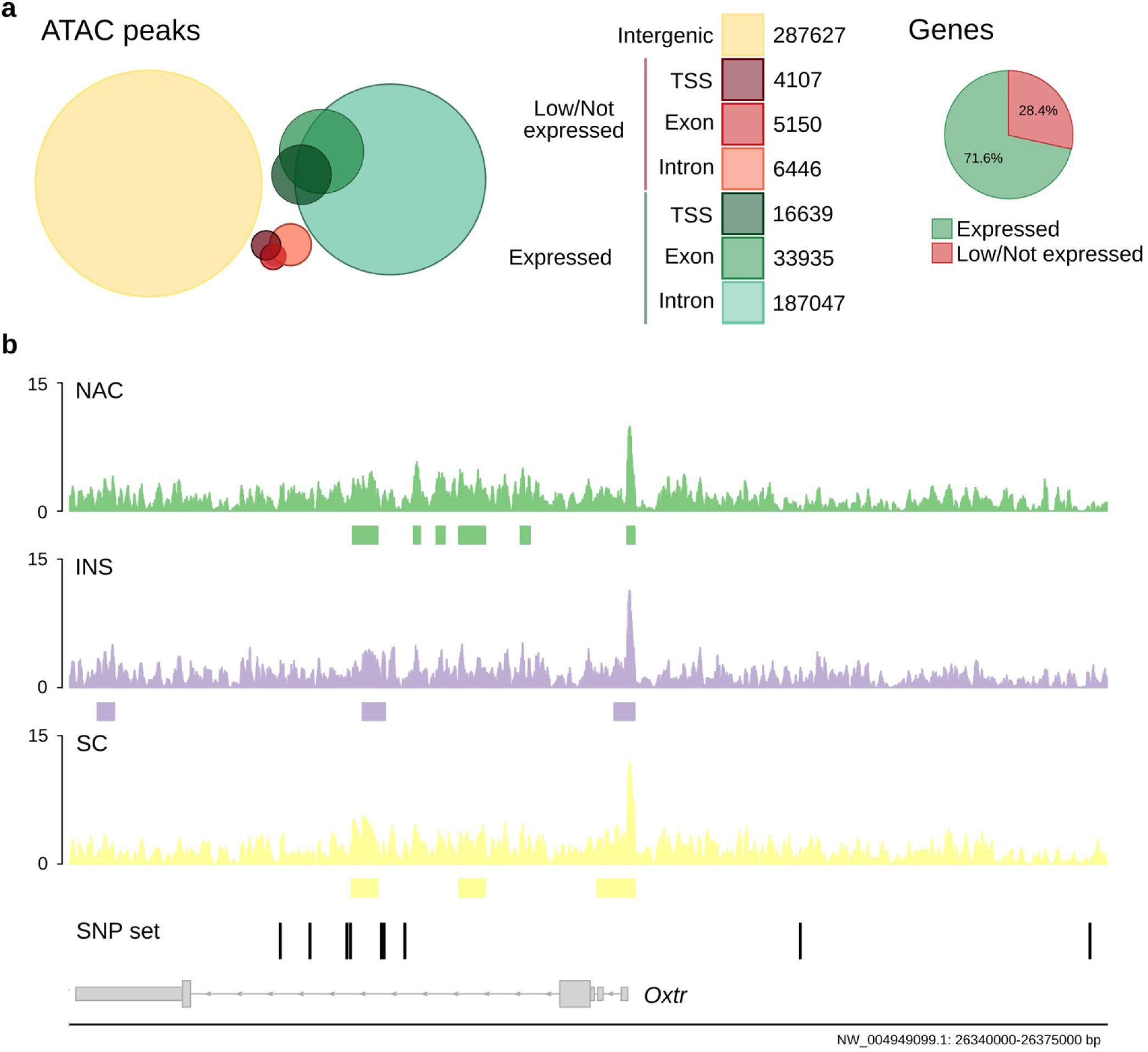
Chromatin accessibility in the *Oxtr* gene locus. **a)** Number of ATAC peaks present in the transcription start site (TSS), exonic, or intronic regions of expressed versus low/not expressed genes, as well as intergenic regions. ATAC peaks preferentially localize to expressed genes rather than low/not expressed genes. **b)** ATAC signal across the prairie vole *Oxtr* gene locus per brain region. Rectangles under the traces indicate significant peaks as identified by Genrich. None of the peaks were significantly different among brain areas or between genotype.

**Supplementary figure 2.**
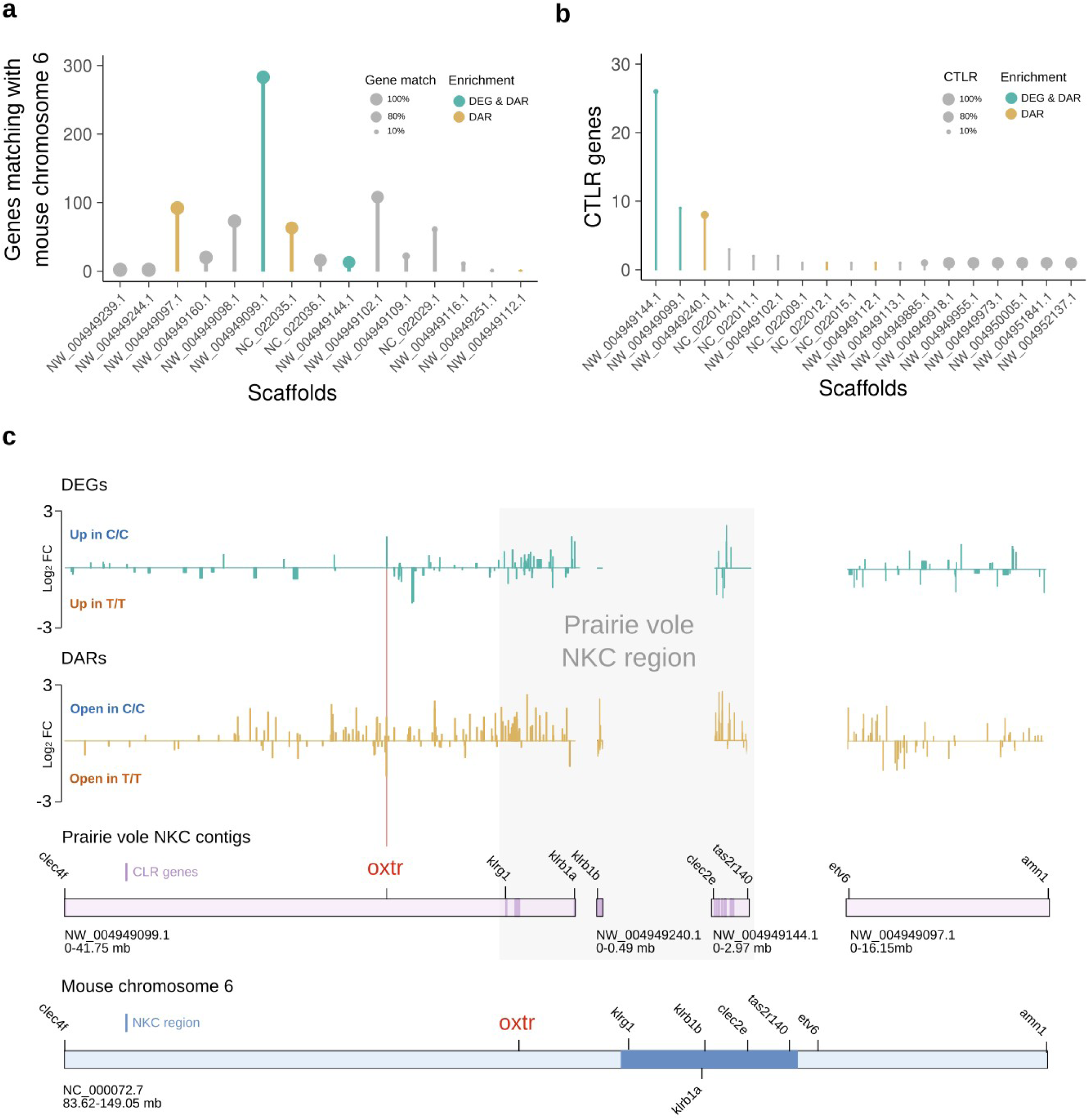
NKC function is divergent between genotypes. **a)** Plot shows the number of annotated genes that a given prairie vole scaffold shares with the mouse *Oxtr* chromosome. scaffolds with no matching genes are not included. Dot size represents the percentage of matching genes. **b)** Plot depicts number of CTLR genes on a given prairie vole scaffold. Scaffolds without CTLRs are not included. Dot size indicates the percentage of genes that are CTLRs. **c)** Upper two plots depict the fold changes of DEGs (green traces) and DARs (gold traces) across NKC scaffolds. Lower plots illustrate how NKC vole scaffolds map to the mouse *Oxtr* chromosome. Scaffolds are on scale, but scales of vole and mouse scaffolds differ.

**Supplementary figure 3.**
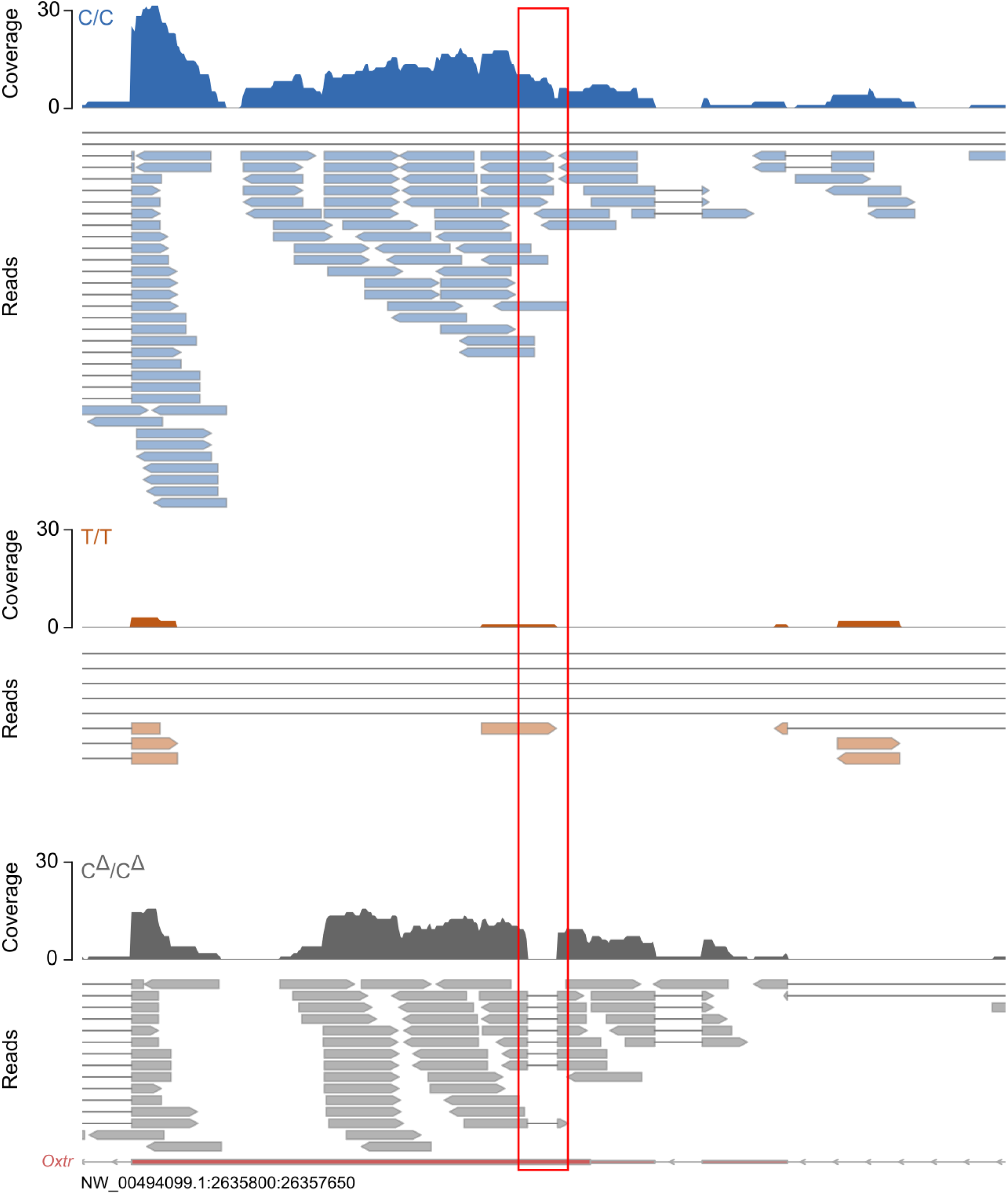
Genetic *Oxtr* deletion is visible in *Oxtr* mRNA reads. Read plots of the 5’-side of the *Oxtr* gene of representative individuals of each genotype. Red box indicates 60bp region in the *Oxtr* coding sequence that is deleted in the systemic OXTR KO but not in wildtype animals.

**Supplementary figure 4.**
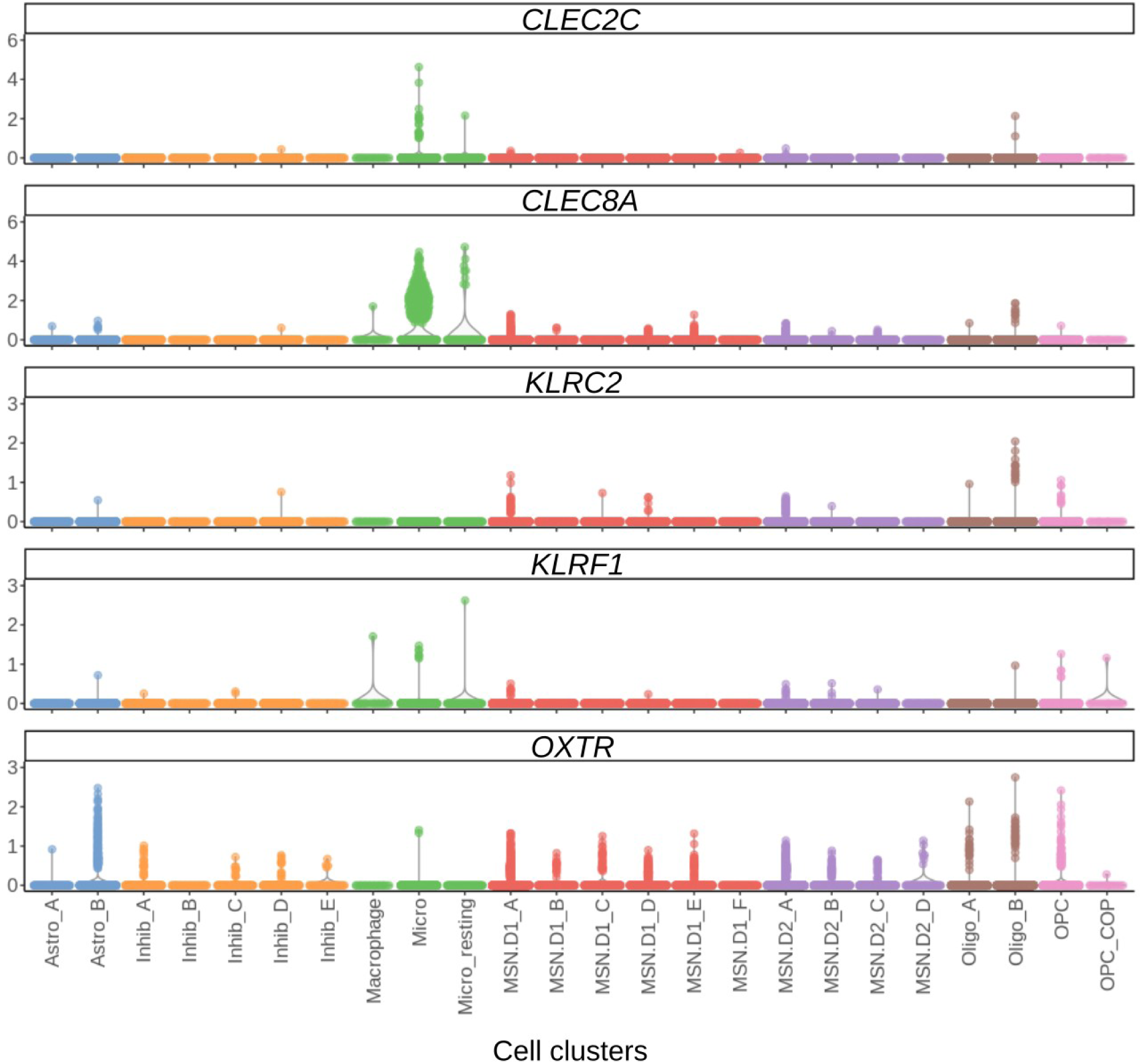
Human NAC expresses *CTLR*s that correlate with *OXTR* abundance. Cell cluster specific expression levels of *OXTR* correlated genes that reside in the NKC as well as *OXTR*.

**Supplementary figure 5.**
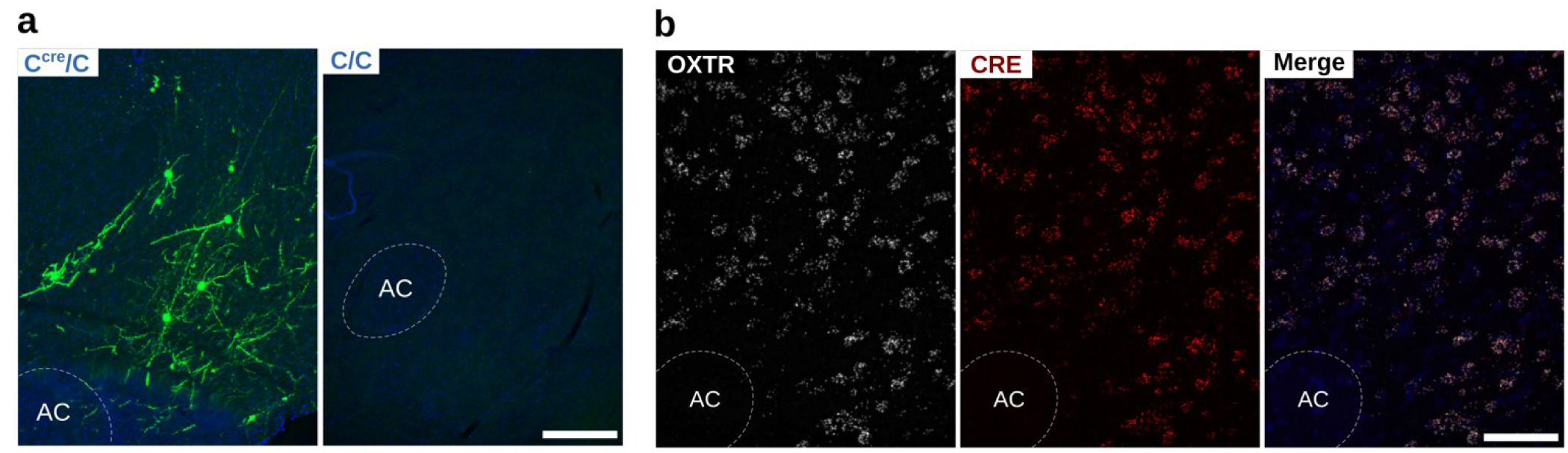
Validation of specific viral-mediated CRE-dependent expression. **a)** Representative images of AAV-injections in the NAC of C^CRE^/C and C/C animals, demonstrating that sparse expression of the fluorophore mGreenLantern is only induced in the presence of a CRE-allele. Scale bar is 100 μm **b)** Representative RNAscope images of the NAC of a C^CRE^/C animal demonstrating that CRE expression is restricted to OXTR-expressing cells. Scale bar is 25 μm.

